# Decoding the Population Activity of Grid Cells for Spatial Localization and Goal-Directed Navigation

**DOI:** 10.1101/021204

**Authors:** Martin Stemmler, Alexander Mathis, Andreas V. M. Herz

**Affiliations:** Bernstein Center for Computational Neuroscience Munich and Department of Biology II Ludwig-Maximilians-Universität München, Grosshadernerstr. 2, 82152 Planegg-Martinsried, Germany; Department of Molecular and Cellular Biology and Center for Brain Science Harvard University 16 Divinity Avenue, Cambridge, MA 02138, U.S.A.

## Abstract

Mammalian grid cells fire whenever an animal crosses the points of an imaginary, hexagonal grid tessellating the environment. Here, we show how animals can localize themselves and navigate by reading-out a simple population vector of grid cell activity across multiple scales, even though this activity is intrinsically stochastic. This theory of dead reckoning explains why grid cells are organized into modules with equal lattice scale and orientation. Computing the homing vector is least error-prone when the ratio of successive grid scales is around 3/2. Silencing intermediate-scale modules should cause systematic errors in navigation, while knocking out the module at the smallest scale will only affect navigational precision. Read-out neurons should behave like goal-vector cells subject to nonlinear gain fields.

---

The regular spatial pattern of grid cell firing (Fig. 1A) is thought to constitute a metric for space (1). For instance, the distance traveled might be estimated by counting the number of episodes in which a grid cell fired. However, the spacing between firing fields, and hence the distance travelled, depends on the movement angle relative to the grid’s orientation (Fig. S1)^1^, which belies the notion of a metric at the single-cell level. The ensemble activity of grid cells could be deciphered to yield a spatial metric (2, 3), but the cipher might need to be learned (4). We show here that a simple, biologically plausible decoder exists.

**Figure 1:**
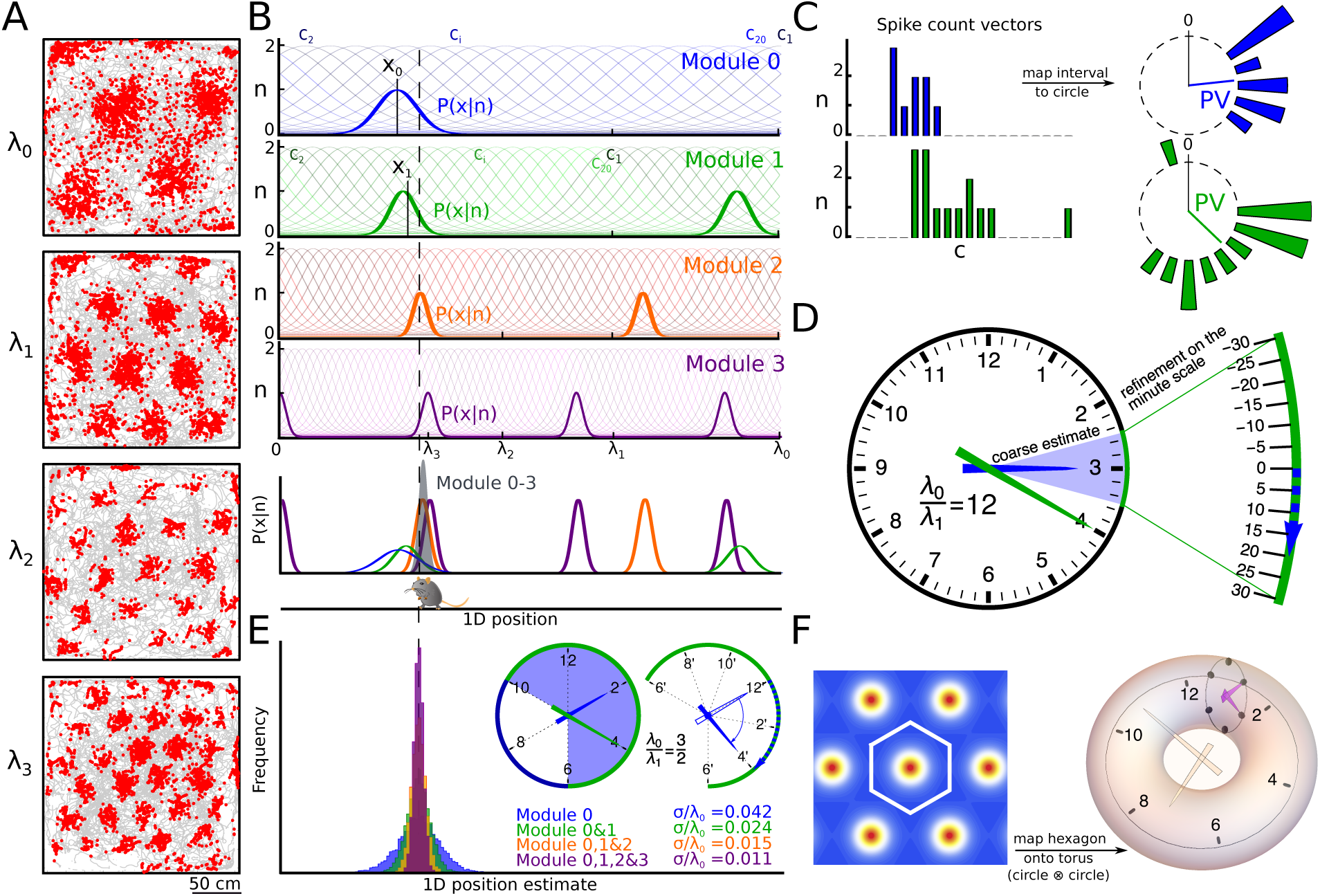
Reading a multi-scale periodic code. (**A**), Grid cells have lattices whose length scales *λ*_*k*_ form a discrete set, ranging from coarse to fine (Figure adapted from Stensola *et al.*, 2012). In gray, the trajectory of a rat in a 2.2 × 2.2 m^2^ enclosure; in red, spikes from 4 grid cells. (**B**), Tuning curves of four nested modules with *M* = 20 grid cells and evenly spaced phases. The animals position yields a spike vector ***n*** in each module. The likelihood *P* (*x ∣* ***n***) at that scale depicts the probability of being at a certain location given the respective spike vector. Modules with smaller spatial periods *λ* have more localized likelihoods, but their multiple peaks result in ambiguous position estimates. The joint likelihood given the responses of all modules, shown in gray, is highly localized and non-periodic. The overall ML estimate is closer to the animal’s position than *x*_0_, the ML estimate of the first module. (**C**), All ML estimates are determined by population vectors (PVs), which are formed by assigning each position *x* to a phase on the unit circle, weighting the number of spikes of each cell by its preferred phase, and then summing, as shown for the first two modules. (**D**), These PVs can be combined for refining the position estimate, similar to how the hour and minute hands of a clock are combined to read the time of the day. In a clock, the ratio of successive scales is 12, as there are 12 hours in each half-day, and in each hour, the minute hand completes one full cycle. (**E**), However, the scale ratio for successive grid modules is not generally integer (the example in (B) has a ratio of 3:2), so at the next scale, a new population vector refines the position estimate by using the earlier estimate *xi* as the center of the range of possible values for *x*_*i+1*_ (Eq. 3). The refined estimate *x*1 in (B) is close though not identical with the ML estimate from this module. Further estimates taking into account module 2 and 3 are recursively calculated (Eq. (S15)). Histograms of these estimates for 2^13^ realizations of the spike vector ***n*** are shown in colors corresponding to the different modules in (B). The relative standard deviations *σ/λ*_0_ highlight that the estimate at each scale successively refines the position estimate (Simulation parameters: *n_max_* = 2, *κ* = 2 & *s* = 3*/*2). (**F**), In two dimensions, the periodicity of the lattice means that the unit cell (white hexagon) can be mapped onto a torus. The position 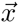 can be read out like a two-dimensional clock with multiple scales.

For each grid cell, the spatial map of firing is captured by a bell-shaped function of space that repeats itself on a hexagonal lattice. All grid cells within one module have the same lattice orientation and grid scale *λ*, but differ in their spatial phase (5, 6). To determine what the neuronal population’s activity reveals about an animal’s location, consider a snapshot of the activity by counting the number of spikes *n*_*j*_ for each neuron in a fixed time window, such that the population’s response is ***n*** = (*n*_1_, *…, n_N_*). When the animal is at position 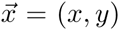 each neuron fires an average of 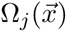 spikes. However, the true number *n*_*j*_ scatters around this value. Assuming Poisson variability and statistically independent neurons, an ideal observer, given the set of spike counts ***n***, will assign the following probability for the animal to be at position 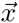:

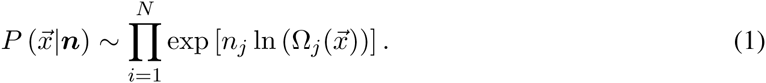

Choosing the most likely position 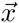 is known as *maximum likelihood* (ML) decoding.

Grid spacings range from about 30 centimeters to several meters (7) in rats. The coarsest scale *λ*_0_ may convey a rough idea of where the animal is, but is subject to some uncertainty *δ*. Eq. 1 predicts generically that an error *δ* introduced at scale *λ*_0_ is *corrected* when a module with scale *λ*_1_ is added; *δ* is reduced to *δ/*[1 + *M*_1_*/*(*M*_0_ *s*^2^)] where *s* = *λ*_0_*/λ*_1_ and *M*_*k*_ is the number of cells at scale *λ*_*k*_. Staggered spatial scales thus implement error correction, and the improvement grows with the number of neurons at the smaller scale (Supp. Mat., Fig. S2).

When space is restricted to one dimension, as on a narrow track (Fig. 1B), ML decoding of a single module reduces to a linear read-out of the population vector average (Fig. 1C), provided the firing rate maps are given by *von Mises* functions (8). These are periodic generalizations of the Gaussian, Ω_*j*_(*x*) = *n*_*max*_ · exp *{κ* [cos(2*π/λ_j_*(*x − c_j_*)) − 1]*}*, where *c*_*j*_ is the preferred spatial phase of cell *j*, and *λ*_*j*_ its spatial period. The population vector (9) points exactly to the most likely position of the animal, and its length conveys the confidence in the position estimate. In analogy to how one tells time using a traditional clock with an hour hand and a minute hand (Fig. 1D), neuronal population vectors lend themselves to an explicit algorithm for decoding position across multiple scales. Assume that the environment fits within the fundamental domain of the module with scale *λ*_0_ (for a generalization, see Supp. Mat.). The population vector is formed by summing over all cells at scale *λ*_0_, weighting each cell’s spike count *n*_*j*_ by its spatial phase *c*_*j*_. The coarse-scale estimate is then

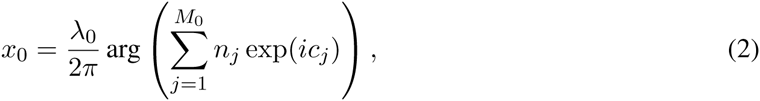

where exp(*ic*_*j*_) = cos(*c*_*j*_) + *i* sin(*c*_*j*_) is a phasor and the arg function computes the angle of a phasor in the complex plane (Fig. 1C). This estimate (Fig. 1B) is refined by the population vector of the second module that contains grid cells with scale *λ*_1_:

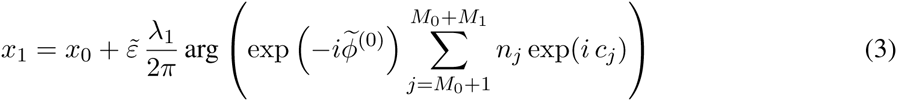

with 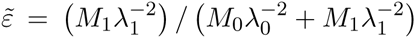 The term involving 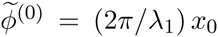 subtracts out the contribution from the earlier position estimate. This procedure can be recursively iterated for all finer modules (Supp. Mat.). Multiple modules thus implement error correction as evidenced by the increasingly more narrow distribution of position estimates (Fig. 1E).

Unlike time, space has more than one dimension. Using three superimposed plane waves as the argument of the *von Mises* function, one obtains a model for hexagonal firing fields in the plane (Fig. 1F). Periodicity in two dimensions means that the lattice’s unit cell is mapped onto a torus. Instead of a single clock, we have two clocks, one for each angle variable on the torus.

In one dimension, a population of neurons with *von Mises* tuning curves yields a posterior probability 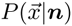 that is also *von Mises*, albeit more peaked. In two dimensions, the lattices of different grid cells may be shifted in spatial phase, as before, or also rotated. Random rotations within a module (Fig. 2A) destroy the hexagonal structure in 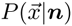 The tolerance to deviations from a perfect alignment can be assessed by computing the grid score (1), which measures the degree of local hexagonal symmetry (Fig. 2B). If the lattice orientations vary by more than 10 degrees, the grid score of 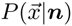 drops precipitously (Fig. 2B). As the lattices of measured grid cells are tightly aligned in orientation (6, 10), the ensemble activity generates a grid-like posterior position probability.

**Figure 2:**
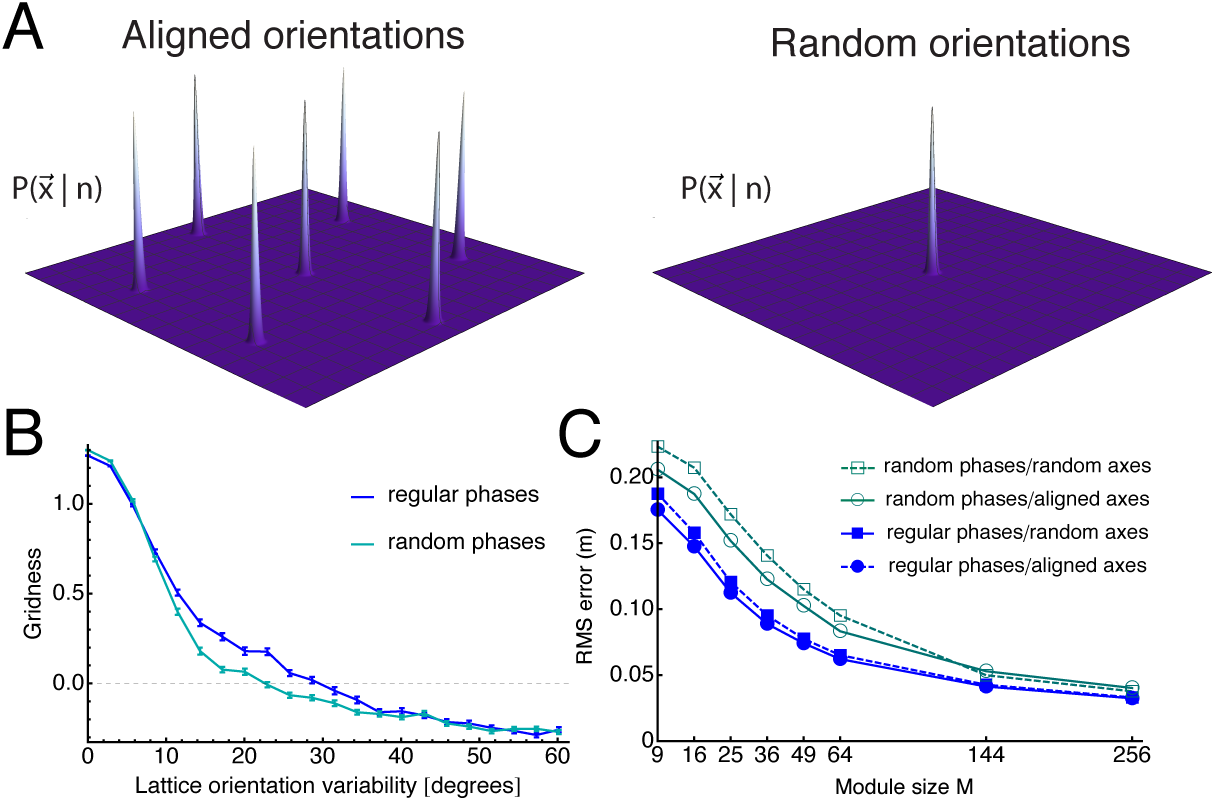
At the population level a periodic representation of 2D space results only for aligned grid lattices. (**A**), 400 neurons with randomly phase shifted but aligned grid-like tuning curves yield a posterior probability distribution 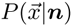 that is hexagonal (left panel). If the lattices are randomly oriented, the hexagonal structure disappears (right). (**B**), The degree of variation in the lattice orientations strongly affects the hexagonal structure in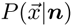, as measured by its ‘gridness’ (1). (**C**), Even for randomly oriented lattices the population response can be decoded by an ideal observer; if the number of neurons is small, aligned lattices result in a lower root mean square (RMS) error. Randomly positioning the lattices, as opposed to evenly spacing them, worsens the error. Size of square box: 1 m^2^.

Whether the lattice orientations are aligned or not, any neural ensemble can be decoded using Eq. (1). For ensembles with few neurons and low peak firing rates, alignment within a single module leads to slightly more accurate position estimates (Fig. 2C). Furthermore, the discrete symmetry axes in 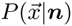 allow the animal to calculate its position by trilateral intersection (Fig. 3A). Three population vectors *μ*_*l*_ are formed by projecting the spatial phase onto the vectors 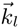 of the hexagonal lattice. The maximum likelihood estimate then reads 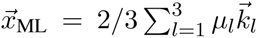. This procedure is equivalent to arranging the population’s spike counts topographically and multiplying by a set of spatial weighting functions that are sinusoidal gratings (Fig. 3B and Supp. Mat.), before forming the population vectors and summing. Implicit in the gratings’ spatial phases is the origin of the coordinate system, yet this origin is arbitrary; for instance, it could represent the animal’s home or a reward location. Switching between locations can be accomplished by rotating the phases (Fig. 3B). The estimate of the homing vector at one spatial scale sets an offset for refining the estimate at the next finer scale (Fig. 3C,S4). Shorter scales imply that the lattice’s tiling of space becomes finer, so this offset resolves the ambiguity associated with the lattice’s periodic nature. The metric read-out of the animal’s position relative to different locations of interest is then the result of a *linear* combination of scales (Fig. 3C). While a population vector code requires the grid axes to be aligned within a module, alignment across modules is not essential. Such an alignment does, however, improve the spatial resolution (Fig. S5) and has been observed experimentally (6, 11, 12).

The total number of grid cells might be as low as 5,000 (13) and downstream neurons might sample from only a small set of cells. To study the limits of grid-cell coding under such adverse conditions, we now consider low firing rates and few neurons per module 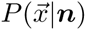 will have a shallow peak, implying that decoding 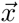 becomes more uncertain and mistakes become more likely. If one reads off the minute hand on a clock, the answer cannot be off by more than thirty minutes; likewise, the worst error in decoding a 2D module with length-scale *λ* is *λ/*2.

**Figure 3:**
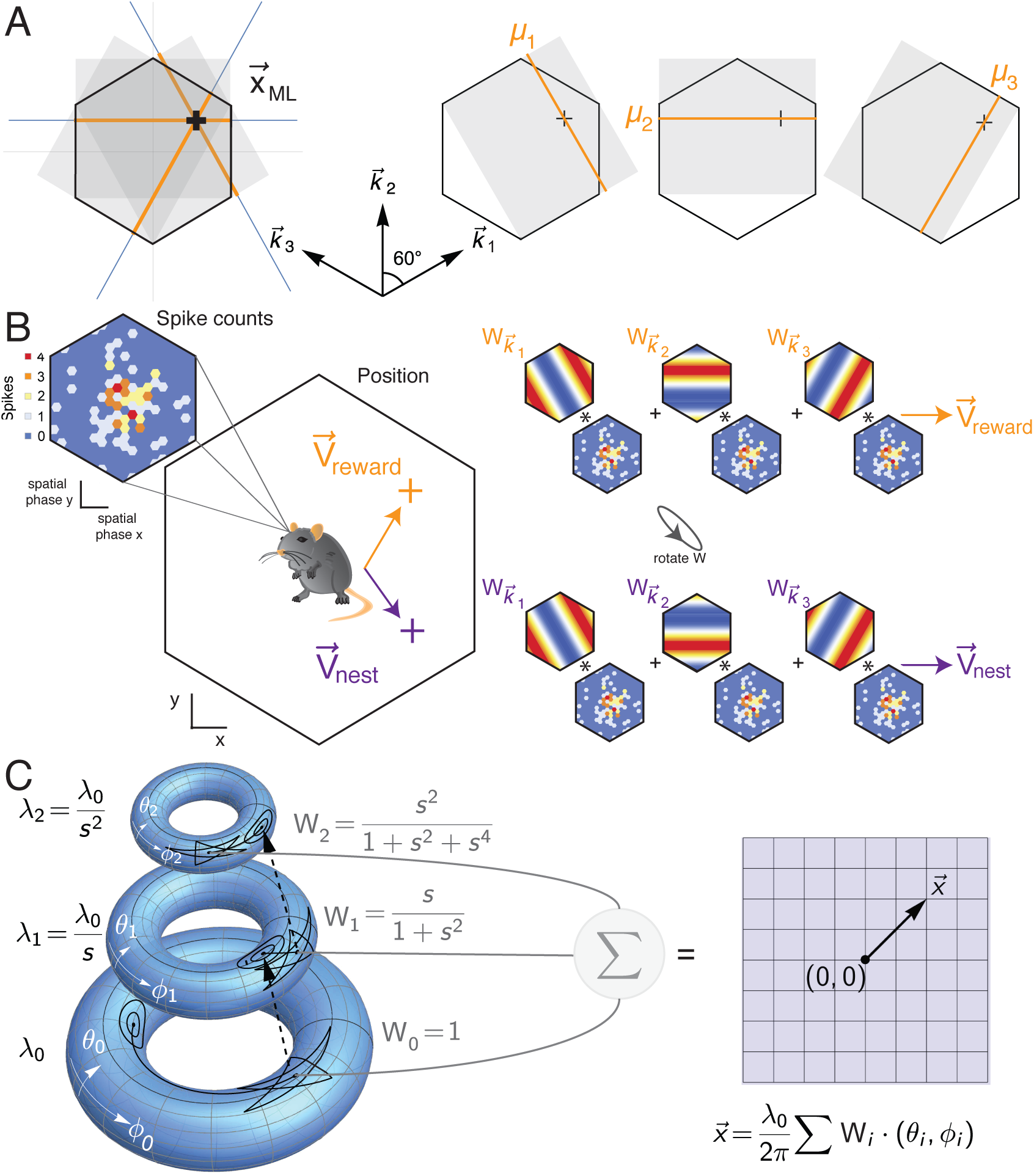
Decoding position in two dimensions. (**A**), The hexagonal lattice has three independent wave vectors, 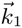, 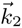, and 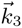, spaced 60° apart. One can transform the hexagonal unit cell into different equally sized rectangles: Form three of these rectangles such that the short edge aligns with the 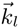’s, Compute the population vector estimate of the position *μ*_*l*_ along the short edge of the rectangle, averaging across the long edge. For each rectangle, this yields a position estimate along the axis 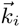, without specifying the position in the orthogonal direction. At the height *μ*_*l*_, draw a line parallel to the long edge in each rectangle. If the projected position estimates were exact, the three resulting lines would meet at one point, the true position of the animal. Otherwise, the three lines form an equilateral triangle, whose center is the ML solution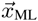,. (**B**), The grid cell population response can yield a homing vector in egocentric coordinates to any point in the environment, such as the location of the nest (purple) or a reward (orange). Rearranging the cells topographically according to spatial phase yields a spike count map. The population vector is formed by multiplying the spike count with cosine gratings, which are aligned along the three axes of the hexagonal lattice. Each such grating is complemented by a weight function phase-shifted by 90° (not shown). The phase of the gratings determines where the homing vector points; rotating the phase of the weights shifts the vector from pointing to the nest to pointing to the reward location. (**C**), Combining population vectors at different scales. At each scale, periodicity implies a map from 2-dimensional Euclidean space onto a torus. The population vectors from the corresponding module yield a vector from the origin (circles) to the estimate of current position (star). This estimate sets the origin of the coordinate system at the next scale. A linear sum of the estimates (*θ_i_, ϕ_i_*) at each length scale, multiplied by weights *W_i_*, produces a precise estimate of the homing vector. These *W*_*i*_’s are functions of the ratio *s* = *λ*_*k*+1_*/λ_k_* of successive length scales. As long as the longest length scale *λ*_0_ is large enough to cover the local environment, the homing vector maps directly back onto Euclidean 2D space.

Refining the position estimate relies on nesting modules at different length scales, with the goal of making the peak in the probability distribution 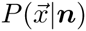 narrower. Figures 4A and 4B illustrate the change in 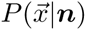 in one spatial dimension upon the addition of a second module with a length scale *λ*_1_. If the scale ratio is *s* = *λ*_0_*/λ*_1_ = 2 (Fig. 4A), the second module increases the probability of the worst possible error: miscomputing the position by *±λ*_0_*/*2.

**Figure 4:**
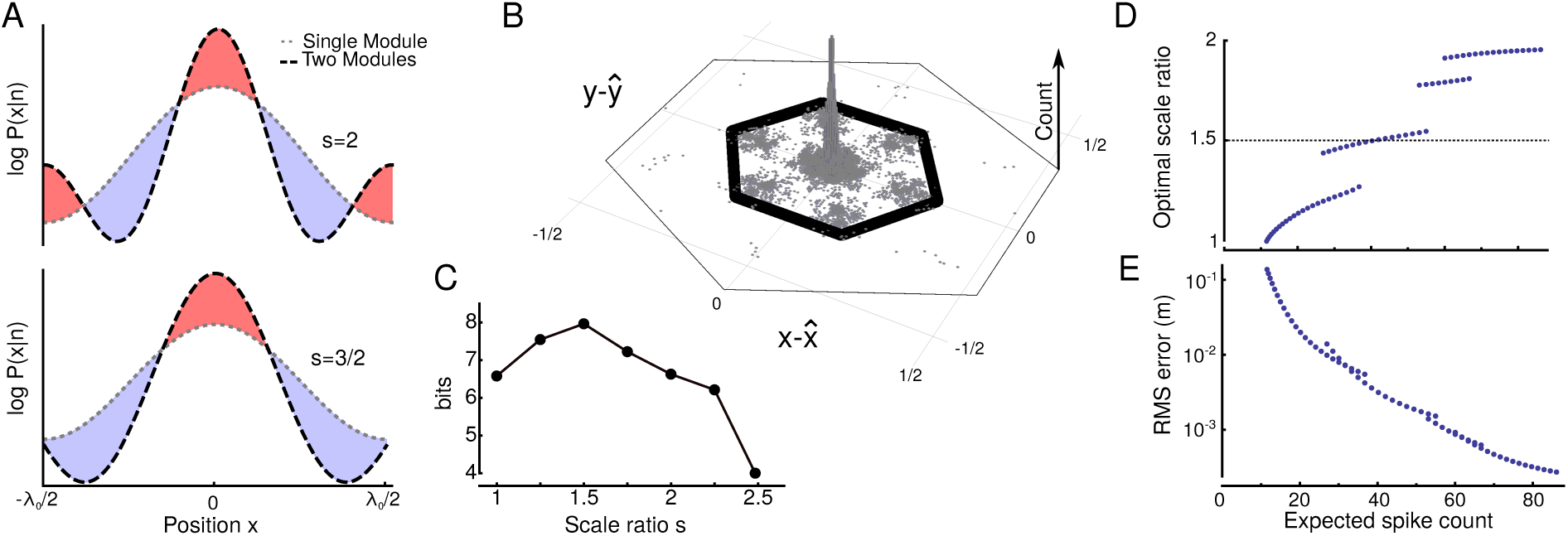
For grid-cell decoding to be robust, the ratio of length scales in successive grid cell modules should be below 3/2. (**A**), Comparison of the log posterior probability log [*P* (*x |* ***n***)] for two scenarios with two modules each but different scale ratios *s* = *λ*_0_*/λ*1 *>* 1. Red shading indicates regions in which the second module increases *P* (*x |* ***n***); blue shading represents a decrease. (**B**), For a grid network composed of four modules with *M* = 64 neurons each, the histogram shows the positions decoded from the spike count relative to the true positions, which were chosen at random 2^15^ times. The length scale of each module was one-half of the next coarser module (*s* = 2), so that the modules interfered. The thin hexagon delineates the spacing between firing fields at the coarsest length scale *λ*_0_, while the thick inscribed hexagon is the unit cell of the lattice. The maximum of the neuron ‘s tuning curve was two; the tuning curve’s shape parameter was *κ* = 2. (**C**), Spatial information in the four-module network as a function of the scale ratio *s*, which reaches a maximum around *s* = 3*/*2. (**D**), The optimal scale ratio *s* depends on the expected number of spikes ⟨***n***⟩ across all four modules and falls into discrete levels. (**E**), The RMS error for the optimal scale ratio *s*, relative to a 1 m^2^ enclosure.

The ratio *s* = 3*/*2, on the other hand, is a safe choice (Fig. 4B). At *x* = *λ*_0_*/*2, the second module contributes a term proportional to cos(*sπ*) to the log-probability; but cos(3*/*2*π*) = 0. Together with the normalization of probability distribution, this trigonometric fact ensures that adding a second module does *not* increase the probability of large decoding errors, irrespective of the number of neurons, the firing rate, or the shape parameter of the neurons’ tuning curve.

The same argument holds when the population code represents two-dimensional space. For the non-ideal ratio of length scales *s* = 2, the histogram of positions decoded from the population spike counts, measured relative to the true position, exhibits a rosetta-like pattern (Fig. 4C): the hexagonal “petals” in this pattern reflect the interference between successive modules.

For low firing rate and small module size *M*, a four-module grid code conveys most information when the length scales obey 1 *≤ s ≤* 2 (Fig. 4D). If the number of bits is *b*, the spike count vector can resolve 2^*b*^ different locations. The greater the number of spikes across the four modules, the higher the scale ratio *s* can be. The optimal *s* falls into discrete levels. The first discrete level distinct from *s* = 1 is centered around *s* = 3*/*2 (dotted line in Fig. 4E).

Multi-scale grid codes can represent vast areas of space (2, 14) or a more limited area with high precision (3). The resolution of such a code could reach a millimeter or less, based on an ideal observer decoding the population response. As we have demonstrated here, the ideal observer is unnecessary: reading the code is both simple and biologically plausible. As in a land survey, measuring the position in two dimensions relies on determining multiple vectors; trigonometry predicts that several neuronal population vectors should be added together to obtain a position estimate. This estimate yields a new, egocentric, vector from the current (allocentric) position 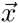to an (arbitrary) origin 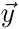; changing the origin of the coordinate system is straightforward, so that a multi-scale grid code could let a foraging animal always know the direction and distance to home. Such a mechanism generalizes population vector-average decoders that have been implicated in visuomotor transformations (9, 15).

While the spatial activity of a single grid cell does not constitute a metric, an ensemble of hierarchically organized grid cells *does* provide a distance measure, as our results show. Even though spatial information is highly distributed across scales, the read-out is a simple linear combination of population vector averages, and implicitly performs error correction. The metric’s accuracy stems from the geometric progression of length scales, ranging from coarse to fine. Such nested scales have been predicted by optimal coding (3). Measured grid cell maps cluster into discrete groups at different length scales, such that the ratio *s* of successive scales lies between 1.4 and 1.7 (5,6). Wei *et al.* (16) derived an optimal scale ratio of *e*^1*/D*^ using a different measure of resolution, where *D* is the dimension of space. In contrast, our argument that *s ≈* 3*/*2 does not depend on *D* (see Supp. Mat.). Because the measured ratio between *λ*_*I*_ and *λ*_*i*+2_ is two (6) or larger (5), silencing an intermediate-scale module should lead to systematic errors (Fig. 4C). On the other hand, removing the grid module with the smallest scale would only affect the fine precision of navigation. Likewise, increasing some grid scales by down-regulating specific cellular conductances (17) should gradually decrease spatial precision according to our theory, whereas it would drastically alter the read-out if grid cells were used for modular arithmetic (2).

Population vector decoding predicts not only that grid cells are grouped into modules, but that lattices within one module should be aligned, as observed experimentally (1, 10). Spatial information would still be present were the lattices randomly aligned or in the absence of modules, but would not be as easily decoded. Yet which neurons read out the grid code? Landmark vector cells in hippocampal area CA1 are one candidate (18, 19), as these cells respond when the homing vector matches a fixed vector describing a specific direction and distance to a landmark. Outside of the hippocampus, circuits in retrosplenial and posterior parietal cortex may be involved, areas essential to memory-dependent spatial navigation (20, 21). A direct link exists between (presumptive) grid cells of presubiculum and retrosplenial cortex (22), whereas the posterior parietal cortex, well-known for the multiplicative interactions between its inputs (23), receives afferents from medial entorhinal cortex (24). One of the key predictions of our theory is that the nervous system will be able to rotate the population vector averages; according to the sum rules of trigonometry, this implies multiplying the read-out weights with cosine-like functions, which could be tested with intracellular in-vivo recordings in downstream areas. Neural mechanisms for such multiplications have been proposed (25, 26), but whether grid cell ensemble activity is decoded in this manner remains a question for further research.

## Footnotes

1 Here and in the following Fig. Sx refers to figure x in the supplementary material.

## Acknowledgements

We thank Edvard Moser for stimulating discussions, Tor Stensola for providing Fig. 1A and Mackenzie Amoroso for graphics advice. Work at the Bernstein Center for Computational Neuroscience Munich was supported by BMBF (01GQ1004A). A.M. received support from DFG grant MA 6176/1-1 and the Marie Curie Fellowship (PIOF-GA-2013-622943 of the European Union’s Seventh Framework Programme FP7 2007-13). Preliminary results have been presented at the Bernstein Conference 2014 (Poster W35), at the Annual Meeting of the Society for Neuroscience 2014 (Abstract 360.05/SS64) and at Cosyne 2015 (Poster III-66 and Workshop II).

